# NR4A3 knockdown ameliorates metabolic dysfunction-associated steatotic liver disease through ATF3 transcriptional repression

**DOI:** 10.64898/2026.06.24.734361

**Authors:** Huiyan Liao, Linlin Zhou, Bo Qin

## Abstract

**Objectives:** The role of nuclear receptor subfamily 4, group A, member 3 (NR4A3) in hepatic steatosis, inflammation, and insulin resistance (IR) within the context of metabolic dysfunction-associated steatotic liver disease (MASLD) remains largely underexplored. Consequently, this study aimed to examine NR4A3’s impact on MASLD and the potential underlying mechanisms.

**Methods:** We aimed to elucidate the functional role of NR4A3 in MASLD through its knockdown in cell culture and animal models. To establish the cell culture model of MASLD, LO2 cells were treated with free fatty acids (FFAs), while male C57BL/6 mice were fed a high-fat diet (HFD) to create the animal model. NR4A3 knockdown was achieved using specific short hairpin RNA (NR4A3-shRNA) in the mice model and three small interfering RNAs (NR4A3-siRNAs) in the cell culture model. The lipids content, fatty acid synthesis, inflammatory factors, and IR were then assessed with and without NR4A3 knockdown. Furthermore, the underlying mechanism through which NR4A3 exerts its influence was explored by analyzing the interaction between NR4A3 and activating transcription factor 3 (ATF3).

**Results:** In the cell culture experiments, the knockdown of NR4A3 significantly decreased the lipids content, fatty acid synthesis, and inflammatory factors in the LO2 cells treated with FFAs in the NR4A3-shRNA group compared with those in the NC-shRNA control group. In the animal model experiments, NR4A3 knockdown in the HFD male C57BL/6 mice significantly ameliorated HFD-induced hepatic steatosis, inflammation, and IR. Mechanistically, the knockdown of NR4A3 downregulated the expression and transcriptional activity of ATF3, resulting in an impaired ATF3 function. ATF3 overexpression significantly reversed lipid accumulation decline and reduced inflammation after NR4A3 knockdown.

**Conclusion:** The downregulation of NR4A3 alleviates MASLD by modulating ATF3, suggesting this may be a promising therapeutic target.

## INTRODUCTION

Metabolic dysfunction-associated steatotic liver disease (MASLD), previously known as non-alcoholic fatty liver disease (NAFLD) ^[1]^, has become a major cause of chronic liver disease worldwide, with an estimated global prevalence of over 30%. It has been reported that 47.3%–63.7% of patients with type 2 diabetes mellitus (T2DM) and 80% of obese patients are affected by MASLD ^[2]^. Given the rising prevalence of metabolic diseases and their frequent association with multi-system disorders, the concept of cardiovascular-renal-hepatic-metabolic (CRHM) syndrome ^[3]^ has been proposed according to the EASL-EASD-EASO guidelines on MASLD ^[4]^, suggesting that MASLD is closely linked to insulin resistance and obesity, which in turn can contribute directly to cardiovascular and kidney damage. MASLD is an increasingly important factor in both the incidence of liver-related and extrahepatic diseases and the mortality from such diseases. Cardiovascular disease and extrahepatic cancer-related issues represent the leading causes of death in patients with MASLD.

The pathogenesis of MASLD is complex, involving disorders of the lipid metabolism, inflammation, insulin resistance (IR), heritability, and the gut microbiome ^[5]^. In particular, the role of lipid metabolism disorders, inflammation, and IR in the pathogenesis of MASLD has been well-documented ^[6]^. MASLD is characterized by a decline in the liver’s capacity to metabolize fatty acids, resulting in excess free fatty acids (FFAs) in the body, which can undergo *de novo* lipogenesis (DNL) and cause an accumulation of toxic lipid species in hepatocytes, resulting in lipotoxic metabolites. These metabolites can induce endoplasmic reticulum (ER) stress and hepatocellular injury, activating the immune system and promoting the release of inflammatory cytokines from innate immune cells ^[7]^. Additionally, MASLD is often associated with IR, as lipid accumulation can occur with IR and hyperinsulinemia ^[8]^. In particular, IR in the liver and peripheral tissues can exacerbate a dysregulated lipid metabolism, promoting hepatic injury and inflammation ^[9]^. This inflammation, in turn, can exacerbate hepatic and systemic IR by impairing insulin signaling pathways, further promoting DNL and creating a vicious cycle that drives disease progression ^[10]^.

Nuclear receptor subfamily 4, group A, member 3 (NR4A3), also known as neuron-derived orphan receptor 1 (Nor-1) ^[11]^, is part of the orphan nuclear receptor subfamily. It was initially discovered in the nervous system but is now recognized for its widespread expression in various tissues, particularly those with high metabolic activity, such as the brain, heart, skeletal muscle, adipose tissue, kidneys and liver ^[12]^. In a previous study using the dataset GSE89632 ^[13]^ from the GEO database, we found that NR4A3 was significantly upregulated in liver samples from MASLD patients. Current knowledge suggests that NR4A3 can activate multiple target genes, including CDKN2A interacting protein (CDKN2AIP) ^[14]^, growth-associated protein 43 (GAP43) ^[15]^, phospho1 ^[16]^, P65 ^[17]^, aldolase A (ALDOA), and phosphofructokinase-1 liver type (PFKL). As a transcription factor, NR4A3 plays a critical role in various cells. Zhu et al. recently showed that the knockdown of NR4A3 mitigates acute pancreatitis-induced injury by reducing oxidative stress, mitochondrial damage, and apoptosis ^[18]^. Additionally, Close et al. reported that NR4A3 knockout mice showed an increase in β-cell mass and improved glucose tolerance ^[19]^. These findings suggest that NR4A3 may play an important role in inflammation and glucose metabolism. Nevertheless, the function of NR4A3 in hepatocytes, especially within the context of MASLD, has yet to be elucidated and remains largely unknown. Consequently, building on our preliminary findings and previous studies in the literature, this research aimed to elucidate the functional role of NR4A3 in MASLD through its knockdown in cell culture and animal models.

Activating transcription factor 3 (ATF3) is a transcription factor induced by cellular stress, is crucial for regulating metabolic and immune processes ^[20]^. Kim et al. showed that ATF3 was highly expressed in the livers of Zucker diabetic fatty (ZDF) rats and in patients with MASLD and/or T2D ^[21]^, and it has also been reported that the overexpression of ATF3 can reduce fatty acid oxidation through mitochondrial dysfunction, leading to IR and hepatic steatosis ^[21]^. Basak et al. found that preventing ATF3/Tip60 could improve liver function by inhibiting fibrotic remodeling and inflammation in high-fat diet (HFD)-fed mice ^[22]^, while Fang et al. reported that IRF2BP2 inhibited hepatosteatosis through repressing ATF3 transcription ^[23]^. Given these findings, we also aimed to investigate ATF3’s role in MASLD development and specifically its association with NR4A3.

In the present study, we found that NR4A3 knockdown ameliorated HFD-induced hepatic steatosis, inflammation, and IR associated with hepatosteatosis in MASLD in both *in vivo* and *in vitro* models of MASLD. Furthermore, we investigated the downstream mechanisms of NR4A3 in the development of MASLD, and notably found that ATF3 was involved in the pathological development of MASLD as one of the downstream targets of NR4A3. These new findings suggest this may be a potential new target for MASLD therapy.

## MATERIALS AND METHODS

### Reagents and antibodies

Palmitic acid (PA) and oleic acid (OA) were purchased from Sigma (St. Louis, MO, USA). The assay kits for assessing the total cholesterol (TC), triglyceride (TG), alanine transaminase (ALT), and aspartate aminotransferase (AST) were purchased from Nanjing Jiancheng Bioengineering Institute (Nanjing, China). The mouse insulin enzyme-linked immunosorbent kit was purchased from ELK Biotechnology (Tianjin, China).

Antibodies against NR4A3 were purchased from Abways (Shanghai, China). Antibodies against ATF3 and phosphorylated forkhead box protein O1 (p-FOXO1) were purchased from Affbiotech (Jiangsu, China). Antibodies against phosphorylated insulin receptor substrate 1 (p-IRS1), protein kinase B (AKT), p-AKT, glycogen synthase kinase 3 (GSK3β), p-GSK3β, p-P38, extracellular signal-regulated kinase (ERK), p-ERK, c-Jun N-terminal kinase (JNK), p-JNK, nuclear factor of kappa light polypeptide gene enhancer in B-cells inhibitor-alpha (IkB-α), p-IkB-α, and p-P65 were purchased from CST (Danvers, USA). Antibodies against IRS1 and P38 were purchased from Abcam (Cambridge, UK). Antibodies against FOXO1 and P65 were purchased from Proteintech Group, Inc (Wuhan, China). β-actin was purchased from TDYbio (Beijing, China). Secondary antibodies were purchased from ASPEN (Wuhan, China).

### Mice

Male C57BL/6 mice (19–21g) were obtained from SiPeiFu Biotechnology (Beijing, China). All the animal experiments were approved by the Institutional Animal Care and Use Committee (IACUC) of the Bestcell Model Biological Center (IACUC Issue No.2024-07-09C). The mice were initially fed via one-week adaptive feeding, after which the mice were randomized into four weight-matched groups (n=5 per group), namely: the normal control (NC), HFD, HFD plus non-targeting short hairpin RNA (shRNA) control (HFD + sh-NC), and HFD plus NR4A3 knockdown (HFD + sh-NR4A3) groups. The mice in the NC group were fed a standard chow diet for 12 weeks, while the mice in the HFD, HFD + sh-NC, and HFD + sh-NR4A3 groups were fed HFD (20% protein, 60% fat, 20% carbohydrate, D12492 formula) for the same duration. Immediately prior to the dietary intervention with HFD, the mice in the HFD + sh-NR4A3 group received a tail vein injection of AAV8 particles (Fenghui, China) containing shRNA plasmids for targeting NR4A3, while those in the HFD + sh-NC group were given an injection of a non-targeting shRNA control. Following the 12-week period, the mice were euthanized, and liver tissues and serum samples were then collected for subsequent analysis.

### Metabolic assays and serum biochemistry in the mice

The mice body weight and blood glucose levels were monitored, and glucose/insulin tolerance tests (GTT/ITT) were performed at specific time points. The serum levels of ALT, AST, TC, and TG were quantified using an automatic biochemical analyzer. All the assays were performed according to the manufacturers’ protocols.

### Pathological examinations

The paraffin-embedded liver tissue sections were subjected to pathological examination using various histological techniques and via light microscopy. Specifically, hematoxylin and eosin (H&E) staining was used to assess the general histology and any pathological changes in the samples. Periodic acid–Schiff (PAS) staining was performed to evaluate carbohydrate metabolism and tissue composition, while to assess lipid accumulation and steatosis, Oil Red O (ORO) staining was performed on frozen liver sections to visualize the neutral lipids to aid the evaluation of steatosis and other lipid-related pathologies.

### Immunofluorescence staining

The paraffin-embedded liver sections were also subjected to immunofluorescence staining to visualize the target antigens/proteins. Briefly, the sections were incubated with primary antibodies overnight at 4 °C, followed by one-hour (h) incubation at room temperature (RT) with fluorophore-conjugated secondary antibodies. Images were then captured using a fluorescence microscope to analyze the localization and distribution of the target proteins within the liver tissue samples.

### Real-time quantitative polymerase chain reaction (qRT-PCR)

Total RNA was extracted from the liver tissues or cells using TRIpure Total RNA Extraction Reagent (ELK, China). Reverse transcription was carried out using EntiLink™ 1st Strand cDNA Synthesis Super Mix (ELK, China). Quantitative PCR was conducted using EnTurbo™ SYBR Green PCR SuperMix (ELK, China) on a QuantStudio 6 Flex Real-Time PCR system (Thermo Fisher Scientific, USA). Primers were purchased from Invitrogen (Shanghai, China), with the sequences listed in Supplementary Table 1. The gene expression levels were quantified relative to β-actin mRNA, which served as the internal control.

### Western blot analysis

For the Western blot (WB) analysis, the liver tissue samples were first homogenized and lysed in RIPA buffer supplemented with protease and phosphatase inhibitors. The protein concentration of the lysates was determined using a BCA assay to ensure equal loading. Equivalent amounts of protein were loaded onto SDS-PAGE gels and transferred to polyvinylidene fluoride (PVDF) membranes. After blocking with 5% non-fat milk to prevent non-specific binding, the membranes were probed with primary antibodies overnight at 4 °C. Following at least three washing steps, the membranes were incubated with secondary antibodies conjugated to horseradish peroxidase (HRP). All the antibodies, along with their respective dilutions and catalog numbers, are listed in Supplementary Table 2. AlphaEaseFC 4.0 software was used to analyze the results for the WB semi-quantitative analysis.

### Adeno-associated virus (AAV) delivery to the liver

AAV8 particles (ELK, China) were obtained containing shRNA plasmids targeting NR4A3, as well as non-targeting control sequences. The mice were given a tail vein injection of 100 μL of the virus (1 × 10^12^ viral genomes) to ultimately achieve AAV8-mediated NR4A3 knockdown. Liver tissue samples were collected 12 weeks after injection, and the efficiency of virus transduction was evaluated by qPCR.

### Cell culture and treatment

The LO2 cell line was purchased from iCell (Shanghai, China). The cell culture media for the LO2 cells consisted of RPMI-1640 medium supplemented with 10% heat-inactivated fetal bovine serum (FBS) and 1% penicillin-streptomycin (P/S). The LO2 cells were treated with 1 mM FFAs, specifically a mixture of OA and PA in a 2:1 ratio, for 24 h to establish an *in vitro* model of MASLD ^[24]^.

### Cell transfection

The LO2 cells were seeded in 6-well plates the day before transfection and were transfected when the confluency reached 30%–50%. Each well was transfected with 5 μL of transfection reagent Lipofectamine 2000 (Thermo Fisher Scientific, USA) and 5 μL of diluted small interfering RNA (siRNA) (ELK, China). After 6 h, the transfection medium was replaced with complete medium, and the cells were then cultured for an additional 48 h. The NR4A3-targeting siRNA expression system included a non-specific control siRNA (siRNA-NC) and three NR4A3-targeting siRNAs, with the following sequences:

siRNA-NC: 5’-GAACAGGAUAGUGAGGUCCCUTT-3’;

siRNA-NR4A3-1,5’-GCAGACAUACAGCUCGGAAUATT-3’;

siRNA- NR4A3-2,5’-GCUGAGCAUGUGCAACAAUUCTT-3’;

siRNA-NR4A3-3: 5’-AGAAGAUCAGACAUUACUUAUTT-3’.

### Dual Luciferase Reporter Activity Assay

Wild-type and mutant DNA fragments covering the ATF3 promoter (Fenghui, China) were inserted into the pGL3-Basic reporter backbone. These constructs were co-delivered with specific siRNAs into HEK293T cells via transfection. Forty-eight hours after transfection, cellular luciferase activity was quantified with the Dual Luciferase Reporter Gene Assay Kit (Vazyme, China) following the manufacturer’s standard protocols.

### Statistical analysis

SPSS version 26.0 statistical software was used to analyze all the data in this study. All the experimental data were obtained from three completely independent repeated experiments and are expressed herein as the mean ± SD. Comparisons between the two groups were performed using a t-test. Multi-group data were analyzed using one-way analysis of variance (ANOVA). The significance level was set at α= 0.05, with p < 0.05 considered statistically significant.

## RESULTS

### NR4A3 expression was upregulated in FFA-treated hepatocytes in vitro and in fatty livers in vivo

To determine whether NR4A3 is involved in fatty liver disease, we treated LO2 cells with FFAs for 24 h to establish an *in vitro* model of MASLD. ORO staining showed that lipid deposition in the FFA group was significantly higher than in the control group (**Figs 1A&B**), while qPCR analysis showed that the expression of NR4A3 was significantly elevated in the FFA group (**Fig. 1C**). Similarly, in the *in vivo* model of MASLD, C57BL/6 mice were fed a HFD and compared with the normal control (NC) group. Significant steatosis was observed in the livers of the mice in the HFD group compared with those in the NC group (**Fig. 1D**). Furthermore, ORO staining showed there were more lipid droplets in the HFD group compared to the control group (**Figs 1E&F**), while Western blot analysis revealed that NR4A3 protein expression in the HFD mice was significantly higher than in the NC group. (**Figs 1G&H**). Collectively, these results indicate that the NR4A3 protein levels were upregulated in both human and mouse steatotic hepatocytes, suggesting NR4A3 plays a role in MASLD.

**Figure 1.**
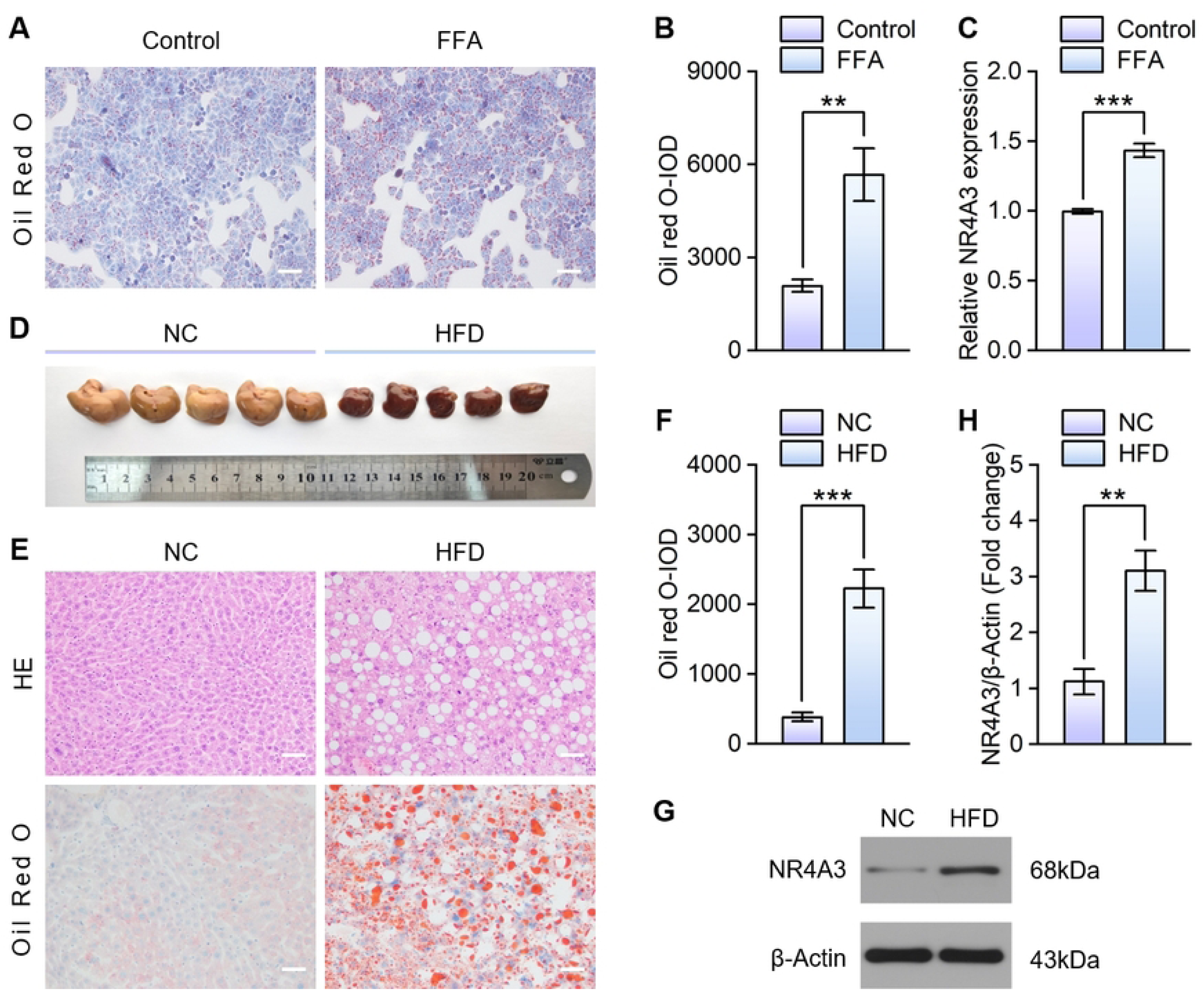
Upregulation of NR4A3 mRNA and protein levels in MASLD. **(A)** Images of ORO-stained LO2 cells from the FFA group and control group (scale bar, 50 μm). The FFA group was treated with a mixture of OA and PA in a 2:1 ratio for 24 h, while the control group received no treatment at 0 h; **(B)** Quantitative analysis of ORO staining; **(C)** Relative NR4A3 expression in LO2 cells detected by qPCR. The mRNA expression levels were normalized to those of β-actin (n = 3); **(D)** Images of liver samples from C57BL/6 mice fed either a normal control (NC) or HFD for 12 weeks (n=5); **(E)** Images of H&E-stained (top) and ORO-stained (bottom) livers from the HFD group and control group (scale bar, 50 μm); **(F)** Quantitative analysis of ORO staining; **(G&H)** Protein expression of NR4A3 detected by Western blot analysis in liver samples from the NC and HFD mice. Data are presented as the mean ± SD. \*\**P* < 0.01, \*\*\**P* < 0.001.

### NR4A3 knockdown reduced FFA-induced lipid accumulation and inflammation in hepatocytes in vitro

To investigate the functional role of NR4A3 in lipid metabolism, we knocked down NR4A3 in LO2 hepatocytes using three NR4A-targeting siRNAs, namely si-NR4A3-1, si-NR4A3-2, and si-NR4A3-3. Western blot analysis confirmed there was a significant reduction of NR4A3 in the three si-NR4A3 groups compared to in the control group (**Figs 2A&B**). Among the three NR4A3-targeting siRNAs, si-NR4A3-2 was selected for the subsequent experiments due to its greatest downregulation. We found that NR4A3 knockdown significantly attenuated FFA-induced lipid accumulation, as evidenced by the ORO staining (**Figs 2C&D**), and decreased the TC and TG levels (**Fig. 2E**). Furthermore, the expression levels of key fatty acid synthesis genes, including acetyl-coA carboxylase alpha (ACACA), fatty acid synthase (FASN), stearoyl-coA desaturase-1 (SCD1), and peroxisome proliferator-activated receptor gamma (PPARγ), were reduced following NR4A3 knockdown (**Fig. 2F**). Given the critical role of chronic inflammation in MASLD, we also evaluated the effect of NR4A3 knockdown on the inflammatory response, and found that the expression levels of inflammatory factors, including tumor necrosis factor-alpha (TNF-ɑ) and interleukin-6 (IL-6), were reduced after NR4A3 knockdown (**Fig. 2G**). Together, these results demonstrate that the knockdown of NR4A3 effectively ameliorated lipid accumulation and inflammation in FFA-stimulated hepatocytes *in vitro*.

**Figure 2.**
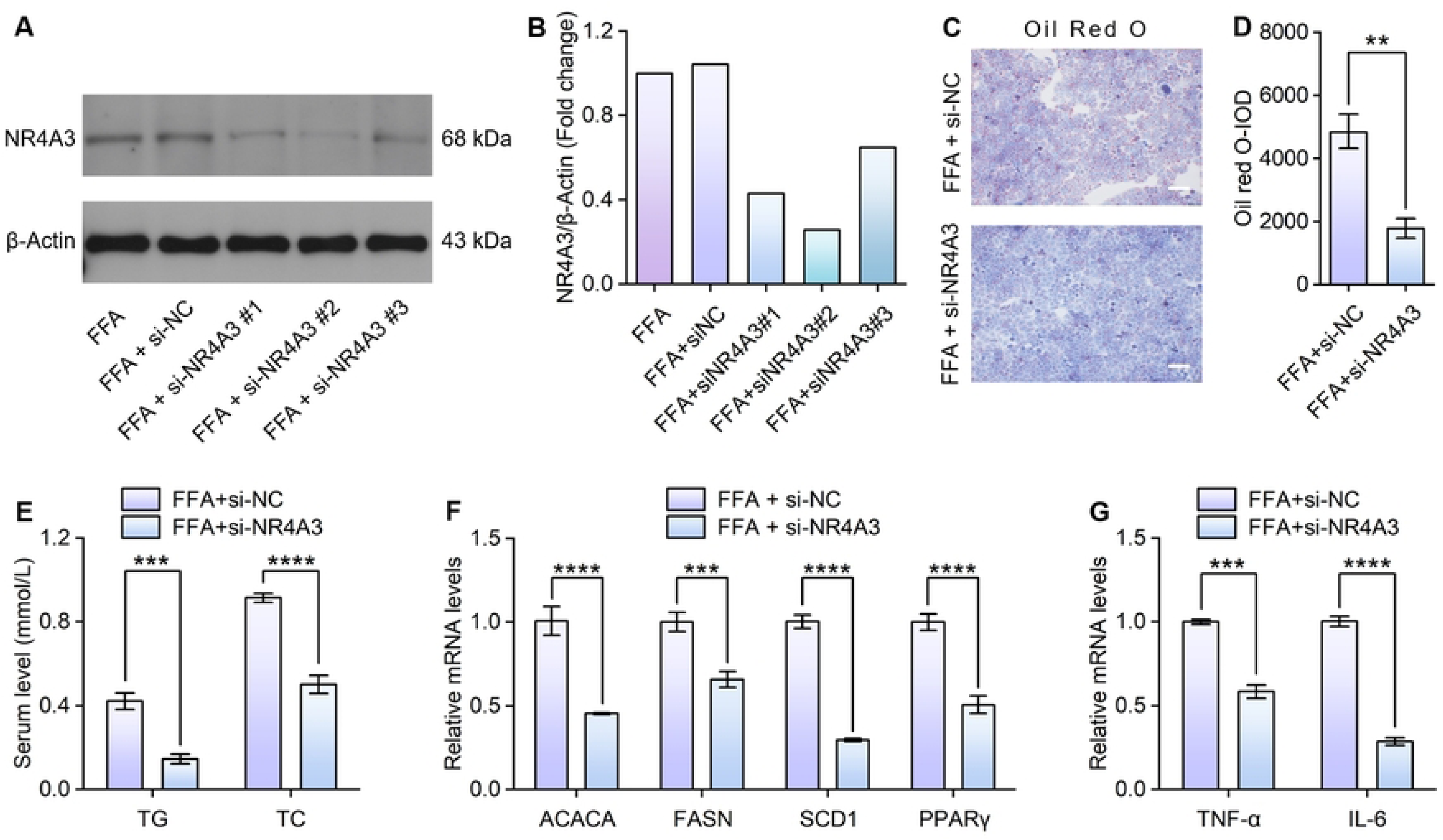
Effects of NR4A3 knockdown on cellular lipid accumulation and inflammation in FFA-treated hepatocytes *in vitro*. (A&B) NR4A3 expression in LO2 cells after exposure to FFAs from different groups determined by Western blot analysis; **(C)** Images of ORO staining from LO2 cells treated with FFAs in the indicated groups (scale bar, 50 μm); **(D)** Quantitative analysis of ORO staining; **(E)** TC and TG contents in LO2 cells treated with FFAs from the indicated groups (n = 3); **(F)** mRNA levels of the fatty acid synthesis in LO2 cells from the indicated groups (n = 3); **(G)** mRNA levels of TNF-ɑ and IL-6 in LO2 cells from the indicated groups,with mRNA expression levels normalized to those of β-actin (n = 3). Data are presented as the mean ± SD. \*\**P* < 0.01, \*\*\**P* < 0.001, \*\*\*\**P* < 0.0001.

### Hepatocyte-specific NR4A3 knockdown mitigated HFD-induced steatosis in the mice

To investigate the functional roles of NR4A3 *in vivo* in lipid accumulation in the liver, we employed gene knockdown techniques in the MASLD mouse model. Through qPCR analysis, we were able to confirm successful hepatocyte-specific NR4A3 knockdown in the livers of the NR4A3 knockdown (sh-NR4A3) mice (**Fig. 3A**). Also, after 12 weeks on the HFD, the NR4A3 knockdown mice exhibited a significant lower weight gain (∼64%) compared with the mice in the control (sh-NC) group (∼98%) (**Fig. 3B**). The liver-to-body weight ratio following 12 weeks on the HFD were also significantly lower in the NR4A3 knockdown mice (**Fig. 3C**). This reduction in liver weight corresponded to a decreased lipid content, as determined by liver lipid measurements (**Fig. 3D**). Consistently, histological analysis using H&E and ORO staining revealed that the NR4A3 knockdown mice had smaller and fewer lipid droplets, suggesting reduced hepatic steatosis (**Figs 3E&F**). We also performed real-time PCR analysis, which revealed that there were lower mRNA levels of key genes in fatty acid synthesis and cluster of differentiation 36 (CD36)-mediated fatty acid uptake in the NR4A3 knockdown mice (**Fig. 3G**), alongside higher levels of genes related to fatty acid β-oxidation, including peroxisome proliferator-activated receptor alpha (PPARα) and carnitine palmitoyltransferase 1 alpha (CPT1α) (**Fig. 3H**). These results collectively indicate that hepatocyte-specific NR4A3 knockdown alleviated HFD-induced liver lipid accumulation.

**Figure 3.**
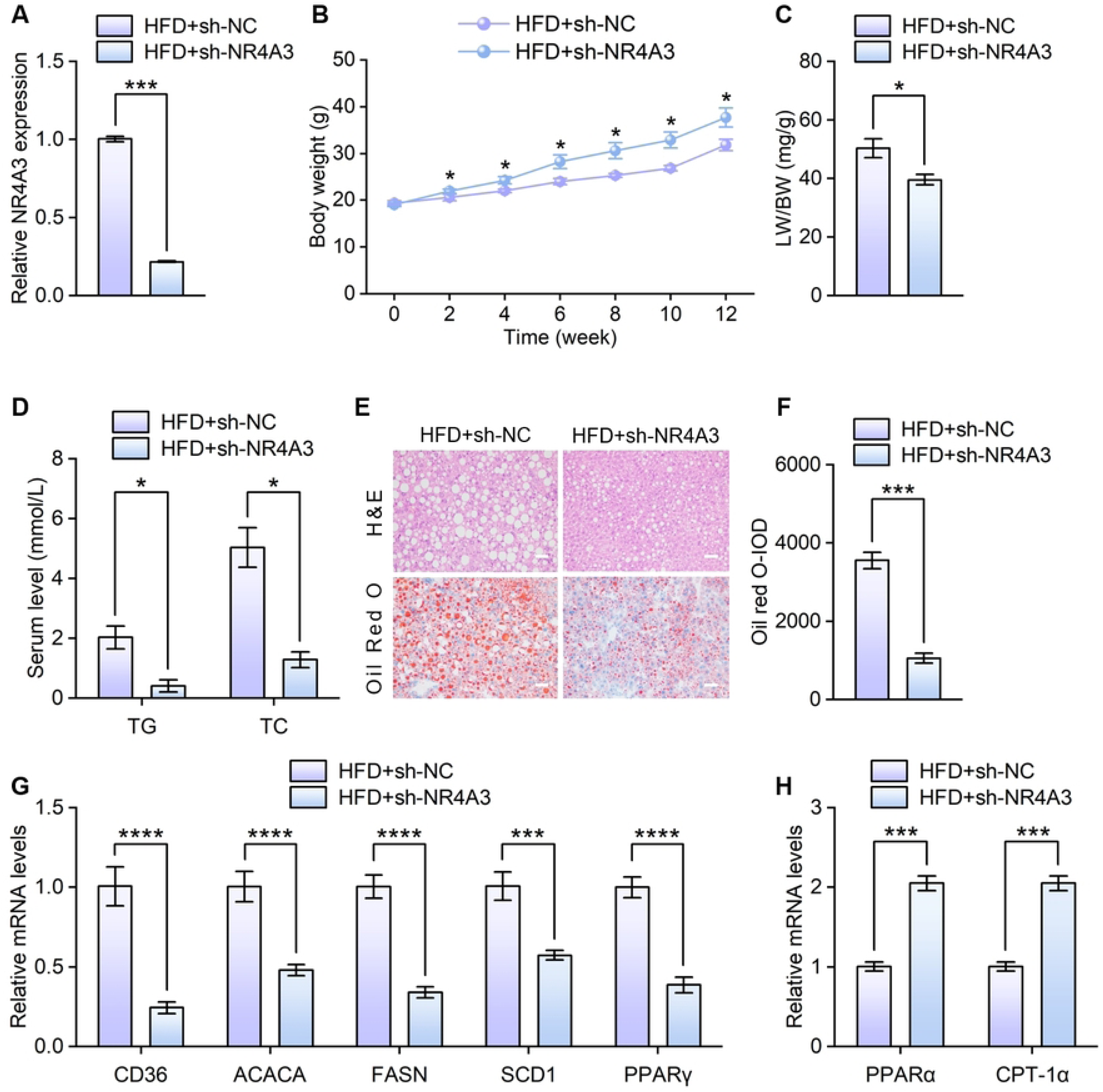
Effects of hepatocyte-specific NR4A3 knockdown on HFD-induced hepatic steatosis in mice. Hepatocyte-specific NR4A3 knockdown mice (sh-NR4A3) and non-targeting control mice (sh-NC) were fed a HFD for 12 weeks. **(A)** NR4A3 mRNA levels in the sh-NR4A3 and control groups; **(B)** Body weight gain curves of the mice in both groups during the 12-week HFD feeding period (n = 5); **(C)** Liver-to-body weight ratio of the mice in both groups (n =5); **(D)** Contents of TG and TC in the livers of the mice in both groups (n=5); **(E&F)** H & E staining (top) and ORO staining (bottom) were used to observe lipid accumulation in liver sections of the mice in both groups (scale bar, 50 μm); **(G)** Relative hepatic mRNA levels of fatty acid uptake and synthesis genes between the two groups,normalized to β-actin (n=3). **(H)** Relative hepatic mRNA levels of fatty acid β-oxidation genes between the two groups, normalized to β-actin (n=3). Data are presented as the mean ± SD. \**P* < 0.05, ***P < 0.001, ****P <0.0001.

### Hepatocyte-specific NR4A3 knockdown mitigated HFD-induced inflammation in the mice

To verify the changes in liver inflammation after NR4A3 knockdown, we assessed the inflammatory indicators in the liver. The results showed that the serum ALT and AST levels (**Fig. 4A**) were significantly reduced in the NR4A3 knockdown mice. Also, the expression levels of the inflammatory factors TNF-ɑ and IL-6 were decreased in the NR4A3 knockdown mice (**Fig. 4B**). Given that the NF-κB pathway ^[25]^ and MAPK signaling have been implicated in the inflammatory response ^[26]^, we investigated the effect of NR4A3 knockdown on the activation of these signaling pathways. Specifically, the mice were fed a HFD for 12 weeks, immunofluorescence images were recorded, which indicated that the nuclear translocation of P65 was decreased in the NR4A3 knockdown group (**Fig. 4C**). Furthermore, Western blot analysis revealed that the phosphorylated active forms p-IkB-α and P-P65 were decreased (**Figs 4D&E**), suggesting that there was a decrease in the NF-κB pathway activity after NR4A3 knockdown. Similarly, decreases in the active forms of the key proteins in the MAPK signaling pathway, namely p-P38, p-ERK, and p-JNK were also observed (**Figs 4D&E**), suggesting that NR4A3 knockdown suppressed the MAPK pathway in the mice. These results indicate that hepatic-specific NR4A3 knockdown ameliorated HFD-induced inflammation.

**Figure 4.**
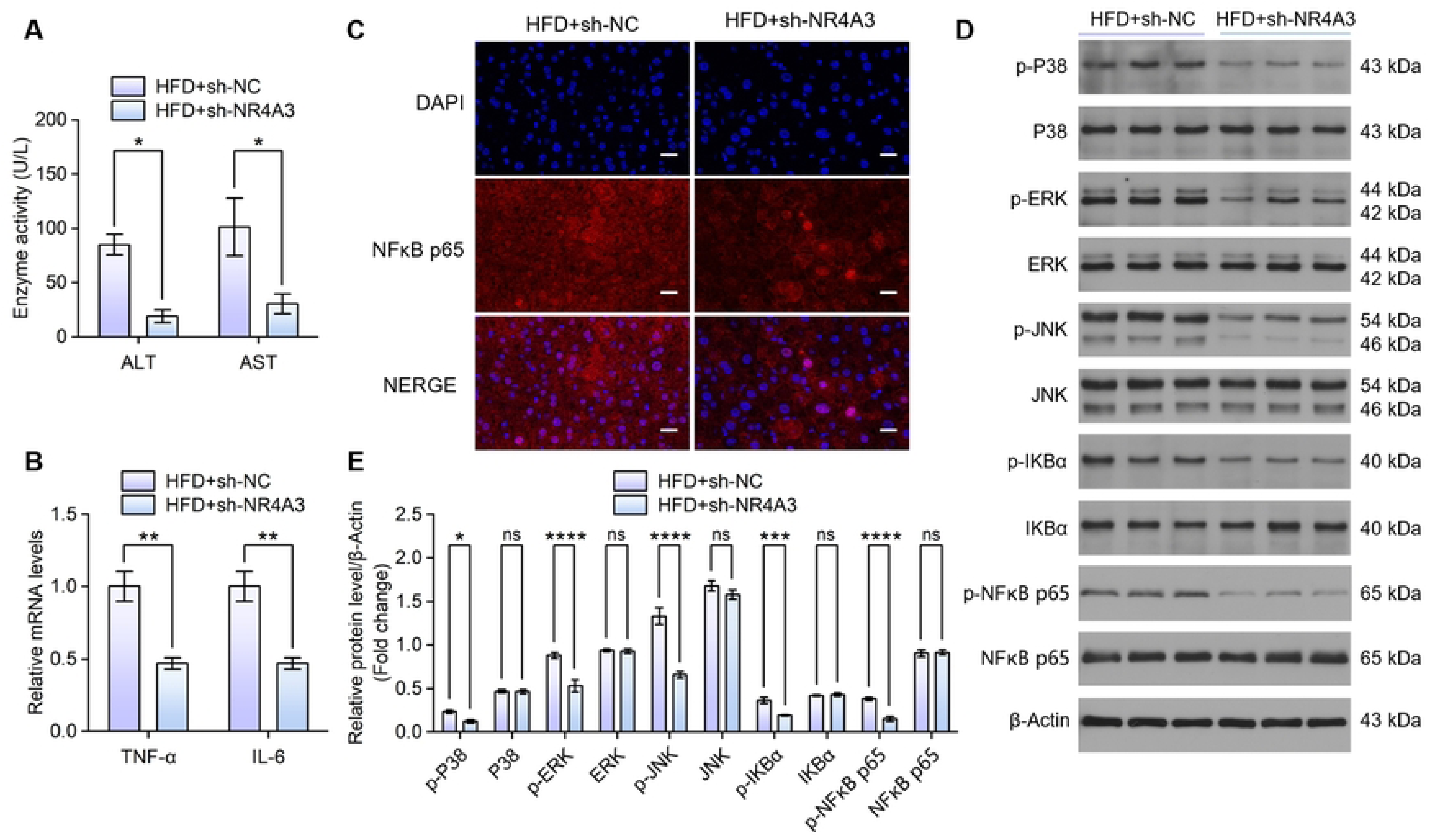
Effects of hepatocyte-specific NR4A3 knockdown on HFD-induced inflammation in mice. sh-NR4A3 and sh-NC mice were fed a HFD for 12 weeks. **(A)** Serum ALT and AST levels in both groups (n = 5); (B) Relative hepatic mRNA levels of the inflammatory cytokines genes between the two groups,normalized to β-actin (n=3); **(C)** Immunofluorescence images showing p65 protein expression in cells (scar bar = 20 μm); **(D&E)** Western blot images and quantitative analysis showing key proteins in the MAPKs and NF-kB signaling pathways in liver tissues from both groups (n =3). Data are presented as the mean ± SD. \**P* < 0.05, \*\**P* < 0.01, \*\*\**P* < 0.001, \*\*\*\**P* < 0.0001. n.s. not significant.

### NR4A3 knockdown ameliorated IR in the HFD-induced mouse model

Considering that MASLD is often associated with IR, as lipid accumulation and inflammation may originate from IR and hyperinsulinemia, we therefore investigated the impact of hepatic NR4A3 knockdown on glucose metabolism in the two groups of mice. The GTT and ITT were performed and revealed that glucose intolerance and IR were ameliorated in the NR4A3 knockdown group, as demonstrated by the significantly smaller AUCs in both the GTT (**Fig. 5A**) and ITT (**Fig. 5B**). During the 12-week period, HFD feeding induced progressive hyperglycemia and hyperinsulinemia in the control mice, while these metabolic disturbances were significantly attenuated in the NR4A3 knockdown mice. Also, the Homeostatic Model Assessment of Insulin Resistance (HOMA-IR) indexes were significantly ameliorated by NR4A3 knockdown (**Fig. 5C**). Consistent with the improved insulin sensitivity, the liver samples from the NR4A3 knockdown group displayed significantly higher glycogen contents (**Fig. 5D**) and lower protein levels of the gluconeogenic enzymes phosphoenolpyruvate carboxykinase (PEPCK) and glucose 6-phosphatase (G6Pase) (**Fig. 5E**). Furthermore, insulin-stimulated activation of the insulin signaling pathway in the NR4A3 knockdown mice was enhanced, as demonstrated by the higher levels of p-IRS1, p-AKT, p-GSK3β and p-FOXO1 in the sh-NR4A3 group (**Figs 5F&G**). These results indicate that hepatic-specific NR4A3 knockdown ameliorated HFD-induced glucose intolerance and IR.

**Figure 5.**
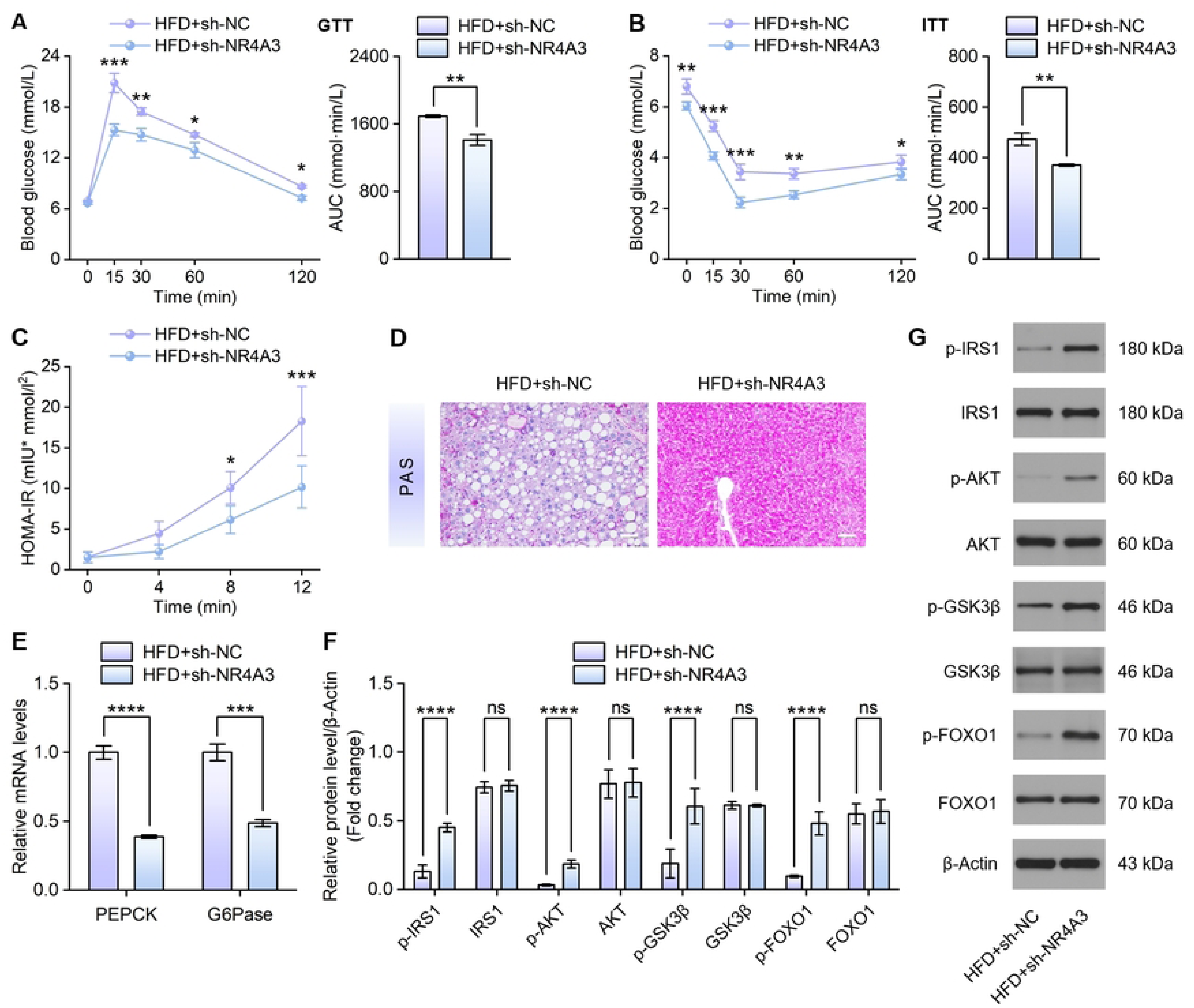
Effects of NR4A3 knockdown on HFD-induced glucose intolerance and the HOMA-IR. sh-NR4A3 and sh-NC mice were fed a HFD for 12 weeks. The GTT **(A)** and ITT **(B)** were performed 12 weeks post-HFD feeding. The AUCs for assessing blood glucose levels were compared between the two groups (n = 5); **(C)** HOMA-IR index curves of the mice in both groups (n =5); **(D)** Representative images of PAS-stained liver sections from both groups (scale bar, 50 μm) (n =3); **(E)** qPCR analysis of hepatic G6Pase and PEPCK mRNA levels in both groups (n =3); **(F&G)** Western blot images and quantitative analysis of hepatic insulin signaling key proteins in both groups, with β-actin serving as a loading control (n =3). Data are presented as the mean ± SD. \**P* < 0.05, \*\**P* < 0.01, \*\*\**P* < 0.001, \*\*\*\**P* < 0.0001. n. s. not significant. AUC, area under the curve.

### ATF3 functions as an immediate downstream molecule of NR4A3 in modulating hepatosteatosis

Accumlating evidence indicates that ATF3 drives liver inflammation through regulating the NF-κB pathway ^[27]^. We therefore proposed that NR4A3 exerts its regulatory effects on hepatic inflammation by targeting ATF3. Herein, we sought to elucidate the mediating role of ATF3 in NR4A3-mediated MASLD pathogenesis. To confirm the regulatory relationship between NR4A3 and ATF3 at the transcriptional level, si-NR4A3 and oe-NR4A3 constructs were applied in cellular experiments. The expression of ATF3 was down-regulated significantly in RT-qPCR after knocking down NR4A3 (**Fig. 6A).** In contrast, expression of NR4A3 induced a robust upregulation of ATF3 transcripts. Collectively, these results indicate a positive regulatory interaction between NR4A3 and ATF3 (**Fig. 6B)**. Next, the JASPAR and NCBI databases were utilized to predict potential binding sites of NR4A3 within the ATF3 promoter. We identified the conserved NR4A3-binding motif **(Fig. 6C)** and subsequently constructed wild-type (WT) and mutant (MUT) ATF3 promoter luciferase reporter plasmids **(Fig. 6D)**. Dual-luciferase assay results confirmed that NR4A3 directly binds to the 5’ UTR region (+1793 to +1809 relative to the transcription start site) of the ATF3 promoter **(Fig. 6E)**. Collectively, these findings demonstrate that NR4A3 directly regulates ATF3 transcription at the promoter level.

**Figure 6.**
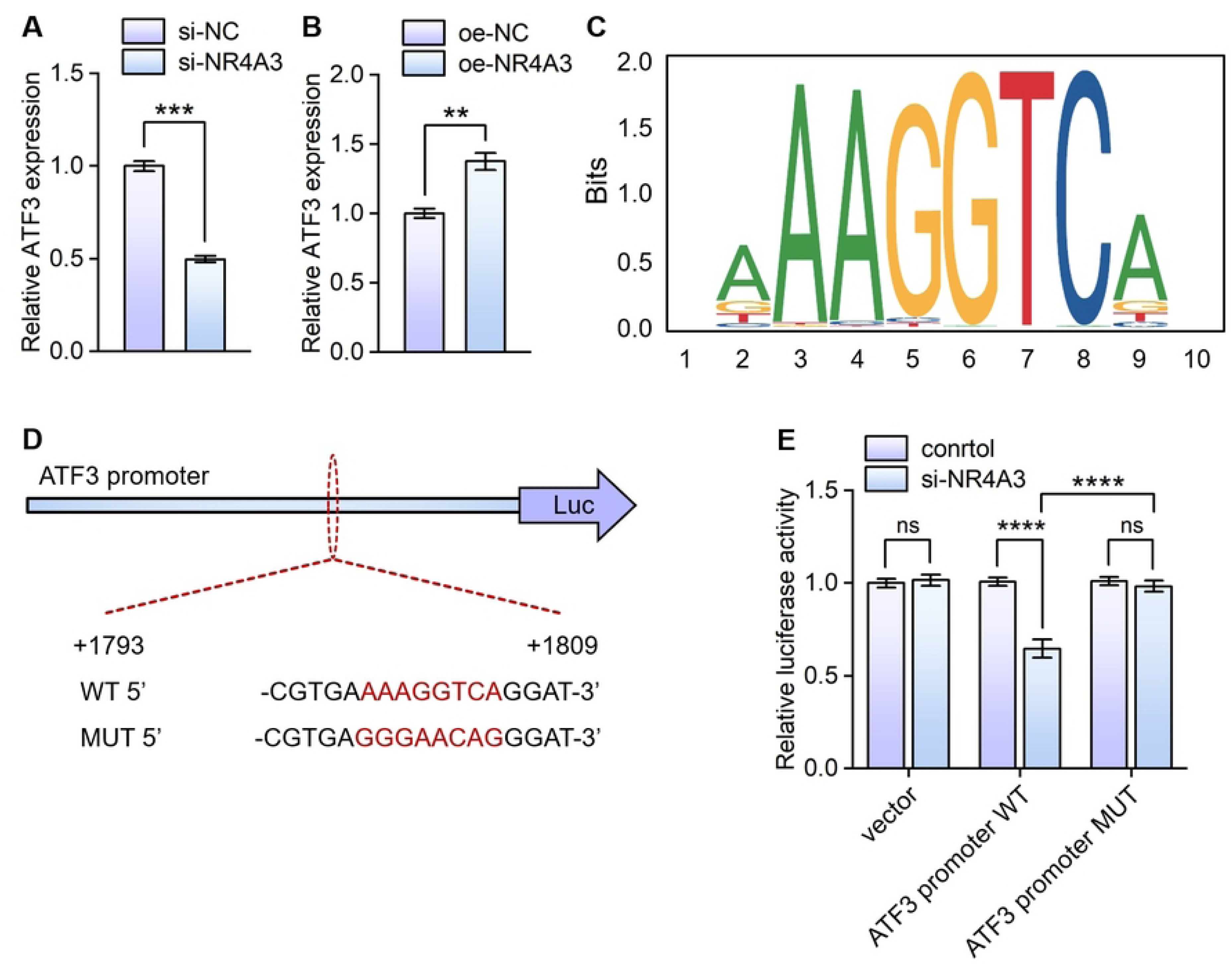
NR4A3 positively regulates ATF3 transcription via direct promoter binding. (A**&**B) qRT-PCR detection of ATF3 mRNA expression after NR4A3 knockdown (A) or overexpression (B) in L02 cells. (C) Sequence logo of the canonical NR4A3 binding motif. (D) Schematic illustration of wild-type (WT) and binding-site mutant (MUT) ATF3 promoter luciferase constructs. (E) Dual-luciferase reporter results showing that knockdown NR4A3 markedly reduces the activity of ATF3-WT promoter, while this effect is abolished after motif mutation. Data are mean ± SD from three independent experiments. **P < 0.01,***P < 0.001,****P < 0.0001.n.s. not significant.

To validate whether ATF3 serves as an essential downstream mediator for NR4A3-induced phenotypic changes, we performed rescue experiments in NR4A3 knockdown LO2 hepatocytes to determine whether ATF3 overexpression could counteract the inhibitory effect on lipid accumulation in NR4A3 knockdown cells (**Fig. 7A**). As illustrated, following treatment with FFAs, significantly less lipid accumulation was observed in the NR4A3 knockdown cells than in the control group. Notably, ATF3 overexpression aggravated lipid accumulation in the NR4A3 knockdown cells (**Figs 7B&C**). Furthermore, we found that ATF3 overexpression significantly reversed the improvement and attenuated the inflammatory response of lipid metabolism genes after NR4A3 knockdown (**Fig. 7D**). These results indicate that the overexpression of ATF3 counteracted the suppressive effects of NR4A3 knockdown on lipid metabolism and inflammatory responses.

**Figure 7.**
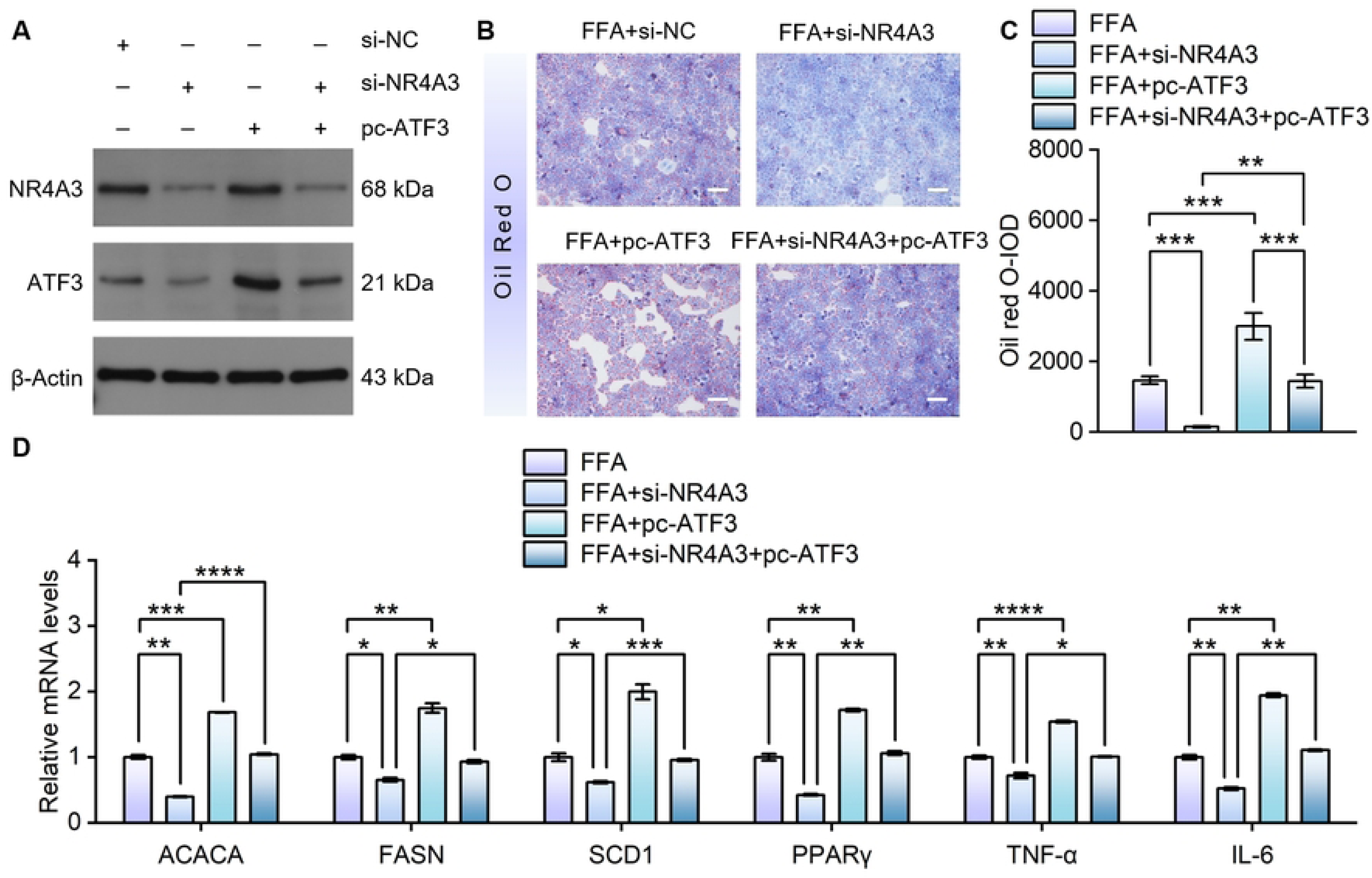
ATF3 overexpression reversed the inhibitory effects of si-NR4A3 on cellular lipogenesis and inflammation. **(A)** Western blot analysis of ATF3 and NR4A3 protein levels in LO2 hepatocytes in four different groups; **(B&C)** Representative images of lentivirus-infected LO2 cells stained with ORO and treated with FFA for 24 h (scale bar, 50 μm). **(D)** Relative mRNA expression levels of fatty acid synthesis and inflammatory cytokines in LO2 cells followed by stimulation with FFAs, with their levels normalized to β-actin; Data are presented as the mean ± SD. \**P* < 0.05, \*\**P* < 0.01, \*\*\**P* < 0.001, \*\*\*\**P* < 0.0001. n.s. not significant.

## DISCUSSION

Early intervention in MASLD can slow the disease progression and reduce the mortality from associated diseases. However, the currently approved therapeutic drugs for MASLD remain limited, with the main options being resmetirom ^[28]^ and the GLP-1 receptor agonist semaglutide ^[4]^. Given that metabolic diseases are characterized by multi-factorial pathogenesis, multi-target drugs are likely to have greater therapeutic potential and consequently a range of multi-target therapies are under investigation. Consequently, there is also an urgent need to identify new therapeutic targets for MASLD. With that in mind, this study investigated the role of NR4A3 in MASLD, a common chronic liver disease with a rising global prevalence, and an urgent need for a better understanding of its pathological mechanisms and the identification of novel therapeutic targets. The major novel findings of this study are summarized as follows: (1) NR4A3 expression was significantly upregulated both in FFA-treated hepatocytes *in vitro* and in fatty livers *in vivo*; (2) NR4A3 knockdown reduced lipid accumulation and inflammation in FFA-treated hepatocytes; (3) Hepatocyte-specific NR4A3 knockdown alleviated HFD-induced steatosis and inflammation in mice; (4) Hepatocyte-specific NR4A3 knockdown improved IR in mice; (5) Mechanistically, the knockdown of NR4A3 downregulated the expression and transcriptional activity of ATF3, resulting in an impaired ATF3 function. ATF3 overexpression significantly reversed lipid accumulation decline and reduced inflammation after NR4A3 knockdown. These findings suggest that NR4A3 may exert its effects through ATF3, which we identified as an immediate downstream target of NR4A3 in modulating hepatosteatosis. Collectively, these findings show that NR4A3 knockdown could alleviate hepatic steatosis and inflammation, as demonstrated in both cellular and animal models, likely by inhibiting ATF3 expression and activity.

Our experimental findings are consistent with the established pathogenic cascade of MASLD. Upon excessive free fatty acid exposure or impaired intracellular fatty acid metabolism in hepatocytes, surplus lipids are converted into lipotoxic intermediates, which trigger endoplasmic reticulum (ER) stress and subsequent hepatocellular damage ^[6]^. Persistent ER stress further activates inflammasome signaling ^[29]^, and the resultant inflammatory response initiates multiple pro-inflammatory cascades, including the NF-kB ^[30][31]^ and MAPK ^[32]^ pathways. As a master transcription factor coordinating metabolic homeostasis and inflammatory responses, NR4A3 may serve as a critical upstream regulator linking lipid toxicity to hepatic inflammation. Prior studies have corroborated the pro-inflammatory function of NR4A3 across multiple disease contexts. Close et al. reported that pro-inflammatory cytokines markedly upregulate NR4A3 expression ^[19]^. In an acute pancreatitis model, Zhu et al. revealed that NR4A3 silencing alleviates oxidative stress, mitochondrial dysfunction and cellular apoptosis ^[18]^. Furthermore, He et al. systematically summarized the broad involvement of NR4A3-dependent inflammation in immune and neurological disorders ^[33]^. Collectively, these studies showed that NR4A3 plays a role in the inflammatory process of various diseases. Consistent with this evidence, in the present study we found that the knockdown of NR4A3 reduces fatty acid uptake and synthesis, promotes fatty acid oxidation, decreases cholesterol deposition, and inhibits protein expression in the NF-kB and MAPK inflammatory pathways, as well as reduces the expression of inflammatory factors such as TNF-α and IL-6, thereby alleviating liver inflammation.

IR can also affect MASLD. In particular, inflammation dysregulates the insulin signaling pathway and contributes to IR, which also further exacerbates metabolic disorder, creating a vicious cycle. Specifically, IR can promote hepatic gluconeogenesis by activating PEPCK and G6Pase through FOXO1. This process not only facilitates MASLD progression but can also cause arteriosclerosis-inducing lipid abnormalities ^[34]^. Close et al. found that NR4A3 expression was significantly upregulated in response to pro-inflammatory cytokines and, to a lesser extent, elevated glucose concentrations in INS and human islet cells; Furthermore, they reported that NR4A3 is upregulated in islets from individuals with T2D, and that the knockdown of NR4A3 can improve glucose tolerance ^[19]^. In another study, Close et al. demonstrated that NR4A3 reduced glucose oxidation, lowered ATP production rates, and inhibited glucose-stimulated insulin secretion ^[35]^. Our findings are consistent with this role in metabolic regulation. In a sh-NR4A3 mouse model under metabolic stress (HFD feeding), we found that the knockdown of NR4A3 improved impaired glucose tolerance and insulin resistance, thereby maintaining glucose homeostasis. This improvement was associated with enhanced insulin signaling pathway activity and a reduction in the expression of gluconeogenic enzymes, which lowered hepatic glucose production. Given that the knockdown of NR4A3 reduced hepatic lipid accumulation and inflammation, improved insulin resistance, and that NR4A3 expression was positively correlated with the development of MASLD, it is considered that targeting NR4A3 represents a promising new therapeutic strategy for MASLD.

In the present study, we also investigated the role of ATF3 and its association with NR4A3 in MASLD. ATF3 is a transcription factor that is known to promote liver inflammation, and has been shown to activate the NF-kB pathway ^[27]^. Kim et al. showed that ATF3 was highly expressed in the livers of individuals with MASLD, and reported that its expression was correlated with hepatic steatosis and disrupted glucose homeostasis ^[21]^. Furthermore, we confirmed that NR4A3 directly regulates ATF3 by binding to its promoter region according to our dual-luciferase reporter assay result. Our rescue experiments provided evidence supporting this hypothesis. Following NR4A3 knockdown, both the mRNA and protein expression levels of ATF3 were found to be significantly decreased. Also, intracellular lipid accumulation was reduced, and the mRNA expression of pro-inflammatory factors such as TNF-αand IL-6 was also significantly downregulated. These findings are consistent with our previous observations that NR4A3 knockdown ameliorated cell lipid deposition and inflammation. However, when ATF3 was simultaneously overexpressed in NR4A3-knockdown cells, the originally suppressed lipid deposition was significantly reversed: the number of intracellular lipid droplets increased, the lipid content rose, and the expression of pro-inflammatory factors was significantly upregulated, nearly returning to the level of the normal control group. Therefore, we concluded that ATF3 was involved in the pathological process of MASLD and acted as a downstream target of NR4A3, indicating that NR4A3 functions in MASLD, at least in part, by suppressing the expression and activity of ATF3.

Despite our novel findings regarding the role of NR4A3 in MASLD, this study has several limitations that should be noted. First, we utilized a single cell line for the *in vitro* experiments; future studies using multiple cell lines and primary human hepatocytes are needed for validation. Second, mechanistically, we confirmed NR4A3 binding to the ATF3 promoter using dual-luciferase assays, while we did not perform co-immunoprecipitation assays to verify protein–protein interaction. The precise molecular interaction pattern between the two molecules therefore remains to be further explored. Third, our initial expression analysis of NR4A3 was only based on public GEO datasets, and we did not validate their expression profiles using clinical human liver biopsy data. In summary, we intend to perform future work, in which we will adopt primary hepatocytes for study, conduct molecular interaction experiments, and collect clinical samples to complement and improve the findings from the current study.

In conclusion, the findings from this study provide initial evidence that NR4A3 plays a novel role in MASLD, and that its knockdown can ameliorate key disease features, including hepatic steatosis, IR, and inflammation. Furthermore, mechanistic investigations indicated that ATF3 may mediate the role of NR4A3 in MASLD. Overall, this study contributes to a better understanding of MASLD and our findings suggest that targeting the NR4A3/ATF3 pathways could offer a promising new therapeutic strategy for the growing population of MASLD patients.

## ACKNOWLEDGEMENT

We thank Medjaden Inc. for their scientific editing of this manuscript.

